# A chemically-defined growth medium to support Lactobacillus – *Acetobacter* community analysis

**DOI:** 10.1101/2021.05.12.443930

**Authors:** Kevin Aumiller, Eric Stevens, Robert Scheffler, Zehra Tüzün Güvener, Emily Tung, Anna B. Grimaldo, Hans K. Carlson, Adam M. Deutschbauer, Michiko E. Taga, Maria L. Marco, William B. Ludington

## Abstract

Lactobacilli and acetobacters are commercially important bacteria that often form communities in natural fermentations, including food preparations, spoilage, and in the digestive tract of *Drosophila melanogaster* fruit flies. Communities of these bacteria are widespread and prolific, despite numerous strain-specific auxotrophies, suggesting they have evolved nutrient interdependencies that regulate their growths. The use of a chemically-defined medium (CDM) supporting the growth of both groups of bacteria would greatly facilitate identification of the precise metabolic interactions between these two groups of bacteria. While numerous such media have been developed that support specific strains of lactobacilli and acetobacters, there has not been a medium formulated to support both genera. We developed such a medium, based on a previous *Lactobacillus* CDM, by modifying the nutrient abundances to improve growth of both groups of bacteria. We further simplified the medium by substituting casamino acids for individual amino acids and the standard Wolfe’s vitamins and mineral stocks for individual vitamins and minerals, resulting in a reduction from 40 to 8 stock solutions. The new CDM and variations of it support robust growth of lactobacilli and acetobacters. We provide the composition and an example of its use to measure nutritional interactions.

## INTRODUCTION

Lactic Acid Bacteria (LAB) and Acetic Acid Bacteria (AAB) coexist in nature in a wide variety of environments (Reese, 1938). These include food fermentations such as wine (Reese, 1938), beer (Dysvik et al., 2020), kefir (da Cruz Pedrozo Miguel et al., 2010; Gulitz et al., 2011), sauerkraut (Wang and Shao, 2018), kimchi (Wang et al., 2016), bread (Li et al., 2021), and cacao beans (Ho et al., 2018). They also have been found together in agricultural feed, such as silage (Guan et al., 2018). LAB and AAB can coexist as well in mammalian and insect gastrointestinal tracts where they have probiotic benefits (Viladomiu et al., 2013), including within the intestine of the genetic model animal, *Drosophila melanogaster*, where lactobacilli and acetobacters are the core types of LAB and AAB respectively (Wong et al., 2011). We note that due to the recent reclassifications within the former *Lactobacillus* genus (Zheng et al., 2020), we refer to the LABs used in this study by their new formal scientific names, *Lactiplanibacillus plantarum* (*Lp. plantarum*) and *Levilactobacillus brevis* (*Ll. brevis*) or collectively by their common name, lactobacillus (plural lactobacilli).

Why LAB and AAB co-occur so prevalently is the subject of ongoing investigations. A synergistic metabolism between lactobacilli and acetobacters has been demonstrated based on sharing of organic short chain fatty acids (SCFAs), including lactic acid and acetic acid. These compounds are generally known to mutually promote growth through cross feeding, such that lactate stimulates acetobacter growth and acetate stimulates lactobacilli (Consuegra et al., 2020; Henriques et al., 2020). There is biomedical relevance of SCFA, including acetate, which are important in human and insect health because their production by the gut microbiome plays a key role in gut epithelial cell metabolism and immune homeostasis (Kim et al., 2018; Rivera-Chávez et al., 2016), potentially underlying the role of LABs as probiotics (Viladomiu et al., 2013). Furthermore, in the mammalian colonic crypts, microbial assemblages include *Lactobacillus* and other Lactobacillales, such as *Streptococcus* and Alphaproteobacteria besides *Acetobacter*, including *Sphingomonas* and *Paracoccus* (Pédron et al., 2012; Saffarian et al., 2019), suggesting there may be a broader phylogenetic pattern of co-existence for these groups, and understanding these relationships in *in vitro* communities could provide insights into their roles in more complex environments with multiple trophic levels such as digestive tract. For instance, cross-feeding may extend beyond SFCAs to other metabolites, including B-vitamins, such as folate and cyanocobalamin, which also impact host health (Degnan et al., 2014; Sannino et al., 2018), and cross-fed nutrients could influence secondary metabolite production (San Roman and Wagner, 2018).

Determining the nutrient requirements for organismal growth (Elli et al., 2000) as well as the metabolites produced (Ponomarova et al., 2017) during fermentation is greatly improved with a defined composition of the growth medium. Chemically-defined growth media have been developed to support lactobacilli growth (Chervaux et al., 2000; Elli et al., 2000; Grobben et al., 1995; Petry et al., 2000; Ricciardi et al., 2015; Saguir and de Nadra, 2007; Savijoki et al., 2006; Wegkamp et al., 2007), yet strain-specific differences in growth requirements between different lactobacilli are widely reported, particularly carbon, amino acid, vitamin, and mineral requirements, (Hayek and Ibrahim, 2013). Construction of a rich CDM could overcome some of these limitations. Furthermore, no media have been developed with the express purpose of supporting lactobacilli*-Acetobacter* communities. To better understand the nutritional dependencies within lactobacillus*-Acetobacter* communities, we developed a rich, chemically-defined medium (CDM) capable of independently supporting growth of both lactobacillus and *Acetobacters* from the fruit fly gut microbiome and other sources in high throughput. We based this medium on the formulation of Savijoki et al 2006. Our medium may be modified to optimize the growth of either lactobacilli or acetobacters to a density of ~10^9^ cells/mL (OD > 1) or to support co-cultures. In this short report, we provide the chemical composition of the medium, some guides on its preparation, an approach to circumvent strain-specific auxotrophies, and some known issues that can result from chemical impurities. We focus our results on the lactobacilli and acetobacters from the *Drosophila* gut microbiome.

## RESULTS AND DISCUSSION

### Formulation of the CDM for lactobacilli growth

With a goal of defining a CDM that supports growth of the lactobacilli and *Acetobacters* isolated from the fruit fly gut (Table S1), we first tested a previous chemically-defined medium that was developed for lactobacillus species (Savijoki et al., 2006). This medium contains ? glucose? All amino acids nucleotides? (*it would be helpful to understand more precisely what was* included) However, that medium supported only limited growth of our strains (OD_600_ of X) and a precipitate formed upon 4 C? room temperature? storage. To prevent precipitation, we modified the medium by reducing the concentration of amino acids from X to X ug/ml. This modified medium supported growth of *Lp. plantarum* LFM1, a *D. melanogaster* isolate, to an OD_600_ of only approximately 0.2 (Table S2), corresponding to ~10^8^ cells/mL. This low value is ~10 fold lower density than reported for the *Lp. plantarum* strain analyzed in the previous study (Savijoki et al., 2006). As strain-specific differences in media preferences are widely reported in lactobacilli (Hayek and Ibrahim, 2013), we further refined the composition by individually adding amino acids, vitamins and nucleotides at 10-fold higher concentrations into the original CDM (Table S3). Higher amounts of tyrosine and cysteine resulted in media precipitation, causing a higher OD_595_ reading, which would interfere with high throughput growth measurements. Improved growth yield with excess alanine and tryptophan was reproducible, thus we increased these concentrations in the CDM to 14 mM and 1.4 mM, respectively (Table 1). We also experimented with increasing or decreasing the total amino acids, vitamins, and nucleic acids (Table S4), which showed we could double these components to increase *Lp. plantarum* growth without substantially affecting other lactobacilli and acetobacters.

**Table 1.** Composition of the chemically 364 defined medium.

| Compound | Concentration | Units | Supplier | Part Number | Stock |
| --- | --- | --- | --- | --- | --- |
| **Base** |  |  |  |  |  |
| MOPS | 40 | mM | Millipore | 475898 | 10x in H<sub>2</sub>O |
| K<sub>2</sub>HPO<sub>4</sub> | 5 | mM | Fisher | P288 | 10x in H<sub>2</sub>O |
| NaCl | 0.2 | mM | Fisher | S271 | 100x in H<sub>2</sub>O |
| NH<sub>4</sub>Cl | 20 | mM | Fisher | A649 | 100x in H<sub>2</sub>O |
| K<sub>2</sub>SO<sub>4</sub> | 10 | mM | Sigma | 746363 | 50x in H<sub>2</sub>O |
| **Metal Salts** |  |  |  |  |  |
| MgCl<sub>2</sub>.6H<sub>2</sub>O | 1 | mM | Fisher | BP214 | 100x in H<sub>2</sub>O |
| MnCl<sub>2</sub>.4H<sub>2</sub>O | 0.05 | mM | Sigma | M3634 | 100x in H<sub>2</sub>O |
| FeSO<sub>4</sub>.7H<sub>2</sub>O | 0.05 | mM | Sigma | F8633 | 100x in H<sub>2</sub>O |
| **Amino acids** |  |  |  |  |  |
| L-Alanine | 14 | mM | Sigma | A7627 | 40x in H<sub>2</sub>O |
| L-Arginine | 0.36 | mM | Sigma | A5131 | 200x in H<sub>2</sub>O |
| Glycine | 3.41 | mM | Sigma | G7126 | 200x in H<sub>2</sub>O |
| L-Lysine | 3.59 | mM | Sigma | L5626 | 200x in H<sub>2</sub>O |
| L-Proline | 1.737 | mM | Sigma | P0380 | 200x in H<sub>2</sub>O |
| L-Histidine | 2.664 | mM | Sigma | H8125 | 200x in H<sub>2</sub>O |
| L-Serine | 7.374 | mM | Sigma | S4500 | 200x in H<sub>2</sub>O |
| L-Threonine | 2.098 | mM | Sigma | T8625 | 200x in H<sub>2</sub>O |
| L-Aspartic acid | 0.083 | mM | Sigma | A9256 | 200x in 1 M HCl |
| L-Asparagine | 4.162 | mM | Sigma | A0884 | 200x in 1 M HCl |
| L-Tyrosine | 1.1035 | mM | Sigma | T3754 | 200x in 1 M NaOH |
| L-Cysteine-HCl | 4.758 | mM | Sigma | C1276 | 200x in H<sub>2</sub>O |
| L-Valine | 4.268 | mM | Sigma | V0500 | 200x in 1 M NaOH |
| L-Glutamic acid | 1.417 | mM | Sigma | G1251 | 200x in 1 M HCl |
| L-Tryptophane | 1.371 | mM | Sigma | T0254 | 200x in 1 M NaOH |
| L-Phenylalanine | 1.513 | mM | Sigma | P2126 | 200x in 1 M HCl |
| L-Glutamine | 3.267 | mM | Sigma | G3126 | 200x in 1 M NaOH |
| L-Leucine | 1.905 | mM | Sigma | L8000 | 2000x in 0.5 M HCl |
| L-Isoleucine | 1.905 | mM | Sigma | I2752 | 2000x in 0.5 M HCl |
| L-Methionine | 0.67 | mM | Sigma | M9625 | 200x in 0.1 M HCl |
| **Nucleotides** |  |  |  |  |  |
| Guanine | 0.033 | mM | Sigma | G6779 | 200x in 0.1 M NaOH |
| Uracil | 0.0445 | mM | Sigma | U1128 | 200x in 0.1 M NaOH |
| Xanthine | 0.0325 | mM | Sigma | X4002 | 200x in 0.1 M NaOH |
| Adenine | 0.037 | mM | Sigma | A2786 | 200x in 0.1 M HCl |
| **Carbon** |  |  |  |  |  |
| Glucose | 1 | % | Sigma | G8270 | 50x in H<sub>2</sub>O |
| Fructose | 0.5 or 1 | % | Sigma | F3510 | 50x in H<sub>2</sub>O |
| Acetate | 0.1 | % | Fisher | S210 | 100x in H<sub>2</sub>O |
| **Optional** |  |  |  |  |  |
| Ascorbic acid | 1.4 | mM | Fisher | AA3623714 | 100x in H<sub>2</sub>O |
| Lipoic acid | 1 | mM | Sigma | T1395 | direct |

We next simplified the medium to reduce the total number of stock solutions. We first replaced the vitamins and minerals with Wolfe’s vitamins and Wolfe’s minerals stocks, which did not noticeably affect growth (not shown). We then replaced the individual amino acids with casamino acids, which are a mixture of purified amino acids derived from peptidase treatment of casein, having a representative abundance of each individual amino acid. This variant of the medium, named CDM_L still required additional cysteine (12 mM final concentration) and tryptophan (2.5 mM final concentration) (Figure S1; Table 2) but supported rapid growth of a variety of *Lp. plantarum* strains from different sources (Table S1) to an OD > 1, corresponding to ~10^9^ cells/mL (Figure 1),.

**Table 2.** Composition of CDM_L 413 (for lactobacilli).

| Stock Solution | Chemical | Final Concentration |
| --- | --- | --- |
| **buffer and salts** |  |  |
| 1 | MOPS | 40 mM (1x) |
| 1 | K<sub>2</sub>HPO<sub>4</sub> | 5 mM (1x) |
| 1 | NH<sub>4</sub>Cl | 20 mM (1x) |
| 1 | Na<sub>2</sub>SO<sub>4</sub> | 10 mM (1x) |
| **metals** |  |  |
| 2 | MgCl<sub>2</sub>*6 H<sub>2</sub>O | 1 mM (1x) |
| 2 | MnCl<sub>2</sub>*4 H<sub>2</sub>O | .05 mM (1x) |
| 2 | FeSO<sub>4</sub>*7 H<sub>2</sub>O | .05 mM (1x) |
| **carbon source** |  |  |
| 3 | glucose / mannitol | 125 mM (1x) |
| **Amino Acids** |  |  |
| 4 | Casamino AAs | 3 g/L |
| 5 | Cysteine-HCl*H<sub>2</sub>O | 1x (0.145 g/L) |
| 6 | Tryptophan | 1x (0.05 g/L) |
| 7 | **Wolfe's Vitamins** | 2x (see composition below) |
| 8 | **Wolfe's Minerals** | 2x (see composition below) |

| Wolfe's Vitamins (100x) | Concentration | units | Supplier | Part Number | Notes |
| --- | --- | --- | --- | --- | --- |
| Ca-(D)-(+)-pantothenate | 0.001 | g/L | Sigma | C8731 |  |
| Nicotinic acid | 0.001 | g/L | Sigma | N4126 |  |
| Para-Aminobenzoic acid | 0.001 | g/L | Nutr. Biochem. | R-238 |  |
| Pyridoxine-HCl | 0.002 | g/L | Alfa aesar | A12041 |  |
| Thiamine HCl | 0.001 | g/L | EMD Millipore | 5871 |  |
| Biotin | 0.0004 | g/L | Sigma | B4639 | in 0.1 M NaOH |
| Folic acid | 0.0004 | g/L | Sigma | F8758 | in 0.1 M NaOH |
| Vitamin B<sub>12</sub> | 0.000002 | g/L | TCI | C0449 |  |
| **Wolfe's Minerals (100x)** |  |  |  |  |  |
| Nitrilotriacetic Acid | 0.3 | g/L | Sigma | N9877 |  |
| MgSO<sub>4</sub>-7H<sub>2</sub>O | 0.6 | g/L | EMD Millipore | MX0070-1 |  |
| MnSO<sub>4</sub>-H<sub>2</sub>O | 0.1 | g/L | Fisher M113 |  |  |
| NaCl | 0.2 | g/L | Fisher S271 |  |  |
| CaCl<sub>2</sub> | 0.02 | g/L | Mallinckrodt | 4160 |  |
| FeSO<sub>4</sub>-7H<sub>2</sub>O | 0.02 | g/L | Sigma F8633 |  |  |
| CoCl<sub>2</sub>-6 H<sub>2</sub>O | 0.02 | g/L | Sigma 202185 |  |  |
| ZnSO<sub>4</sub>-7 H<sub>2</sub>O | 0.02 | g/L | Fisher Z68 |  |  |
| CuSO<sub>4</sub>-5 H<sub>2</sub>O | 0.002 | g/L | VWR 330 |  |  |
| AlK(SO)<sub>4</sub>-12 H<sub>2</sub>O | 0.002 | g/L | Sigma 237086 |  |  |
| H<sub>3</sub>BO<sub>3</sub> | 0.002 | g/L | Fisher A78 |  |  |
| Na<sub>2</sub>MoO<sub>4</sub>-2 H<sub>2</sub>O | 0.002 | g/L | Acros Organics | 446360250 |  |

**Figure 1.**
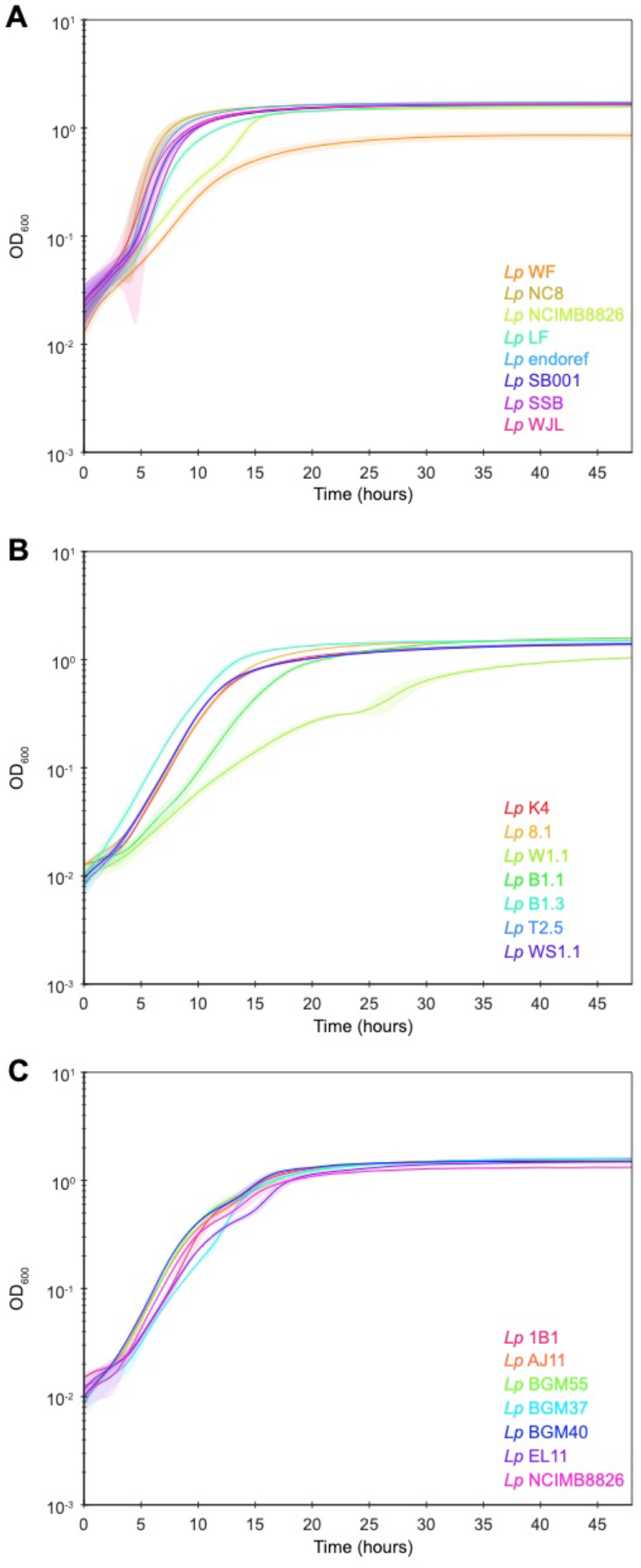
*Lp. plantarum* growth in CDM_L. Most strains grew to OD > 1.5, while *LpWF* grew to OD 0.8. Cultures were inoculated in 96 well microplates from CDM passaged cells at a starting OD of 0.05. N=12 replicates per sample. Solid line is cubic spline-smoothed mean. Shading is s.e.m. Samples in B & C were grown in a separate lab from A, with a separate batches of media and a different plate reader. In panel A, cells were passaged through CDM three times before inoculation for growth measurements. In B and C, cells were inoculated from MRS. Note faster initial growth in A.

We note several technical issues that arose when constructing and testing the various CDMs. First, inocula from nutrient rich agar plates such as MRS agar do not grow well in CDM, but conditioning them by passaging through a mixture of 25:75 MRS:CDM before 100% CDM greatly improved the number of strains that grew. For many strains, the most robust growth resulted after three passages in CDM.

Second, we noticed that when different commercial sources of iron and cobalt were used, a precipitate sometimes formed after 48 hours at room temperature. Re-filtering the media to remove the precipitate did not inhibit *Lp. plantarum* growth, suggesting that lower metal concentrations may be used at least for some strains. As noted earlier, amino acid precipitation was also an issue, which we were able to resolve by reducing the total amount of amino acids (by x-fold) while increasing select ones through the use of casamino acids, cysteine, and tryptophan.

### Modification of the CDM for *Acetobacter* growth

Next, we incubated several strains of *Acetobacter* in CDM_L. *A. pasteurianus* and *A. tropicalis* exhibited extremely limited growth (Table S2). In an attempt to improve growth, we repeated the 10-fold additions experiment used test for improved lactobacilli growth, with ascorbate providing the only growth improvement (Table S3). Because ascorbate caused discoloration of the media after 48 h at room temperature, we only use it optionally. Through a series of trials, we also added lactate, substituted potassium acetate for ammonium acetate, and reduced the amino acid concentrations (by 2.5-fold to formulate a new version of the CDM named CDM_A (Table 3), which supported an OD_595_ of ~1.0 (~10^9^ cells/mL) for the three *Acetobacter* strains tested (Figure 2). We note that it was critical to supply adequate aeration to *Acetobacter* by shaking and poking a hole in the sealing film over the plate. Increased shaking speed increased the clumping of the cultures, which appears as increased variance in the growth curves (Figure 2). Reducing the shaking speed reduced clumping but also reduced the growth rate.

**Table 3.** Composition of CDM_A (for *Acetobacters*).

| Reagent | Concentration | Supplier | Part Number |
| --- | --- | --- | --- |
| **Base** |  |  |  |
| MOPS 40 0.837 8.37 (pH 6.8) | 40 | mM (Millipore) | 475898 |
| K<sub>2</sub>HPO<sub>4</sub> | 5 | mM (Fisher) | P288 |
| NH<sub>4</sub>Cl 18.69 0.1 10 | 20 | mM (Fisher) | A649 |
| K<sub>2</sub>SO<sub>4</sub> 8.28 0.144 7.2 | 10 | mM (Sigma) | 746363 |
| **Metal Salts** |  |  |  |
| MgCl<sub>2</sub>·6H<sub>2</sub>O 0.98 0.02 2 | 1 | mM (Fisher) | BP214 |
| MnCl<sub>2</sub>·4H<sub>2</sub>O 0.05 0.001 0.1 | 0.05 | mM (Sigma) | M3634 |
| FeSO<sub>4</sub>·7H<sub>2</sub>O 0.035 0.001 0.1 | 0.05 | mM (Sigma) | F8633 |
| **Carbon Sources** |  |  |  |
| Glucose | 125 | mM (Sigma) | G8270 |
| Potassium Acetate | 6 | mM (Fisher) | P171 |
| DL-Lactic Acid | 50 | mM (Sigma) | 69785 |
| **Amino acids** |  |  |  |
| Casamino Acids | 6 | g/L BD Bacto | 223050 |
| L-Cysteine HCL | 0.58 | g/L Sigma | C1276 |
| L-Tryptophan | 0.2 | g/L Sigma | T0254 |
| **Wolfe's vitamins 1x** |  |  |  |
| **Wolfe's minerals 1x** |  |  |  |

**Figure 2.**
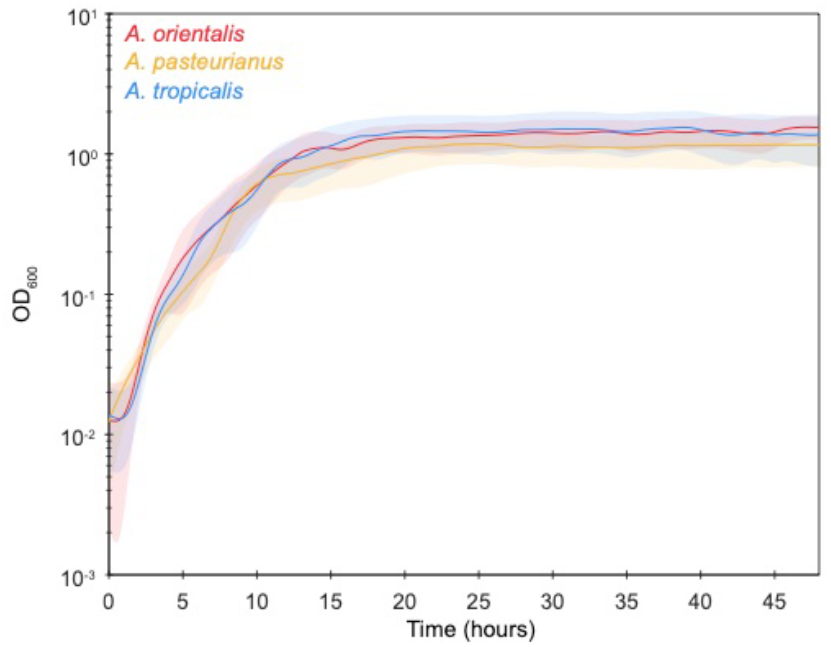
*Acetobacter* strain growth in CDM_A. *A. orientalis*, *A. pasteurianus*, and *A. tropicalis* grow to O.D. >1.0. Cultures were inoculated in 96-well plates from CDM-passaged cells at a starting O.D. of 0.05. N= 12 replicates per sample. Solid line is cubic spline-smoothed mean. Shading is s.e.m.

### An *Ll. brevis* strain shows cross-feeding with an *Acetobacter*

Some preliminary experiments with dropouts of individual nutrients suggested that *Ll. brevis* strain LF, which we isolated from laboratory *D. melanogaster*, had several auxotrophies. This strain grew in Man Rogosa Sharpe (MRS) rich medium but was unable to grow in the complete CDM_L, suggesting metabolic deficiencies (Figure 3A). Growth was restored with the addition of 2.5% v/v MRS to the CDM_L (CDM_L+MRS_2.5 (final OD_600_ = 0.6). One of the auxotrophies appeared to be for folate. We then made CDM_L+MRS_2.5 lacking folate (CDM_L+MRS_2.5-folate). *L. brevis* LF was unable to grow in CDM_L+MRS_2.5-folate, consistent with a folate auxotrophy (Figure 3A).

**Figure 3.**
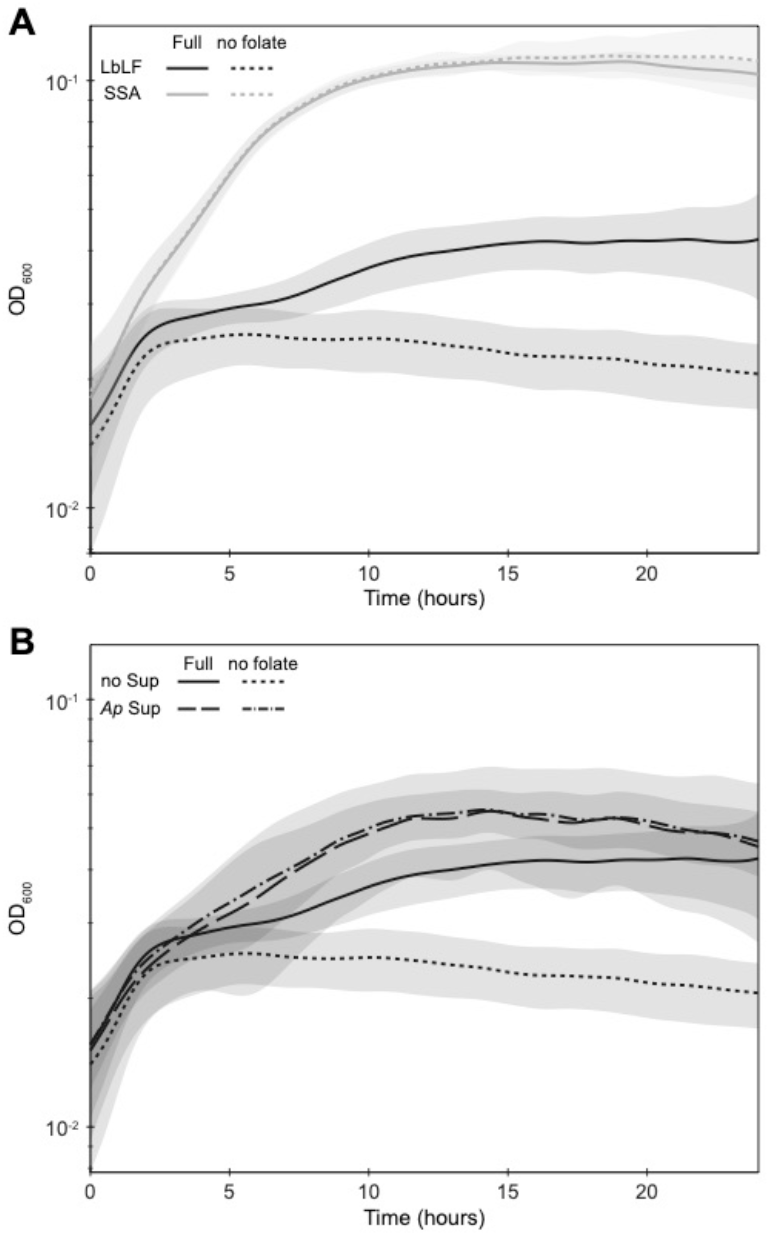
Lactobacillus-Acetobacter co-culture in spent CDM_L+MRS_2.5 that lacks folate. (A) *Levilactobacillus brevis* LF grows in CDM_L supplemented with 2.5% MRS but not when folate is excluded. (B) *Acetobacter pasteurianus*-conditioned media restores *Ll. brevis* LF growth in CDM lacking folate. All samples grown in 96-well plates. N=12 replicates per sample. Solid line is cubic spline-smoothed mean. Shading is s.e.m.

Acetobacters are known B-vitamin producers, with complete folate metabolism (Bernhardt et al., 2019; Sannino et al., 2018). Coculture with *A. pasteurianus* increased *Ll. brevis* LF growth in CDM_L as did a 50:50 mix of CDM_L+MRS_2.5-folate with filter-sterilized spent CDM_L+MRS_2.5-folate from *A. pasteurianus* growth (Figure 3B), indicating that *A. pasteurianus* produces metabolites (likely including folate) that compensate for the *Ll. brevis* auxotrophies. While the overall densities of cells are quite low, this example shows how the base CDM can be used to explore community metabolism in lactobacillus*-Acetobacter* cocultures, even when the medium cannot support the individual strains.

## CONCLUSIONS

The CDM formulated here enables investigation of lactobacillus*-Acetobacter* community metabolism and it may provide a starting point for investigation of more diverse LAB and AAB communities, which are often observed together in nature.

## MATERIALS AND METHODS

### Chemicals

A list of chemicals used in CDM are provided in Table S5.

### Media preparation

Stock solutions were individually prepared in ultra-pure water or in appropriate solvents. pH adjustments were made as indicated in Table 1. All stock solutions were passed through 0.22 μm filter and kept in dark at 4 °C except nucleotides, which were stored at −20 °C. FeSO_4_.7H_2_O was freshly prepared. Medium was initially made with 30 % less water to allow customized additions. Final pH of CDM was adjusted to 6.5 and the medium was passed through a 0.22 μm filter. CDM was stored at 4 °C and used within 2 days.

To make a carbon-free CDM, the following components were combined in the following order in a final volume of 17.5 ml: 8.375 ml ultra-pure water, 2.5 ml MOPS buffer, 0.25 ml K_2_HPO_4_, 0.25 ml NaCl, 0.25 ml NH_4_Cl, 0.5 ml K_2_SO_4_, 0.625 ml L-Alanine, 0.125 ml L-Arginine, 0.125 ml Glycine, 0.125 ml L-Lysine, 0.5 ml L-Proline, 0.125 ml L-Histidine, 0.125 ml L-Serine, 0.125 ml L-Threonine, 0.125 ml L-Aspartic acid, 20 uL of 10 N NaOH for pH correction, 0.125 ml L-Asparagine, 10 uL of 10 N NaOH for pH correction, 0.125 ml L-Tyrosine, 0.125 ml L-Cysteine-HCl, 0.125 ml L-Valine, 0.125 ml L-Glutamic acid, 0.125 ml L-Tryptophane, 0.125 ml L-Phenylalanine, 0.125 ml L-Glutamine, 0.125 ml L-Leucine, 0.125 ml L-Isoleucine, 0.125 ml L-Methionine, 0.125 ml Ca-D-(+)-pantothenate, 0.125 ml Lipoic acid, 0.125 ml Nicotinic acid, 0.125 ml para-Aminobenzoic acid, 0.125 ml Pyridoxine-HCl, 0.125 ml Thiamine-HCl, 0.125 ml Biotin, 0.125 ml Ascorbic acid, 0.125 ml Folic acid, 0.125 ml Guanine, 0.125 ml Uracil, 0.125 ml Xanthine, 0.125 ml Adenine, 0.25 ml MgCl_2_.6H_2_O, 0.25 ml MnCl_2_.4H_2_O, 0.25 ml FeSO_4_.7H_2_O.

Appropriate carbon sources were provided in CDM to grow specific isolates. A final concentration of 1 % glucose and 0.1 % acetate were added for the growth of *L. plantarum*. A final concentration of 1 % glucose, 0.5 % fructose and 0.1 % acetate was optimized for the growth of *L. brevis*. A final concentration of 1 % fructose was used for the growth of *Acetobacter pasteurianus*, and 1 % glucose was used for the growth of *A. tropicalis*.

### Bacterial strains and growth

Bacterial strains used in this study are listed in Table S1. Strains were grown on MRS agar plates at 30 °C incubation for 1 to 2 days. Plate-grown cells were routinely generated from glycerol stocks and plates were used only once. At least 30 colonies [to reduce the odds of a mutant dominating the growth curve] were transferred into 3 ml CDM in a test tube and cultures were shaken at 30°C for 16 to 18 h. Turbidity was adjusted to OD_600_ of 0.5 and used to inoculate media for growth kinetics experiments. Time-course growth experiments were performed in 100 μL CDM in 96-well microplates. After aliquoting 70 μL CDM into each well, indicated carbon sources from sterile stocks were added to appropriate wells and the medium was inoculated with 10 μL culture for a final OD_600_ of 0.05. Plates were sealed with a Breatheasy film. Microplates were incubated in a temperature-controlled plate reader with shaking. OD_595_ measurements were taken every 15 minutes for 24 to 48 hours. M1000 (Tecan) and Magellan (Biotek) plate readers were used for experiments. OD_595_ measurements were background-subtracted using the OD_595_ measurement of medium-only blank controls.

For acetobacter growth, maximal shaking was used, with a speed of 600 RPM at an orbital radius of 3 mm. To improve aeration further, we poked an off-center hole in the Breatheasy film over each well using a fine gauge needle.

---

**Figure S1.**
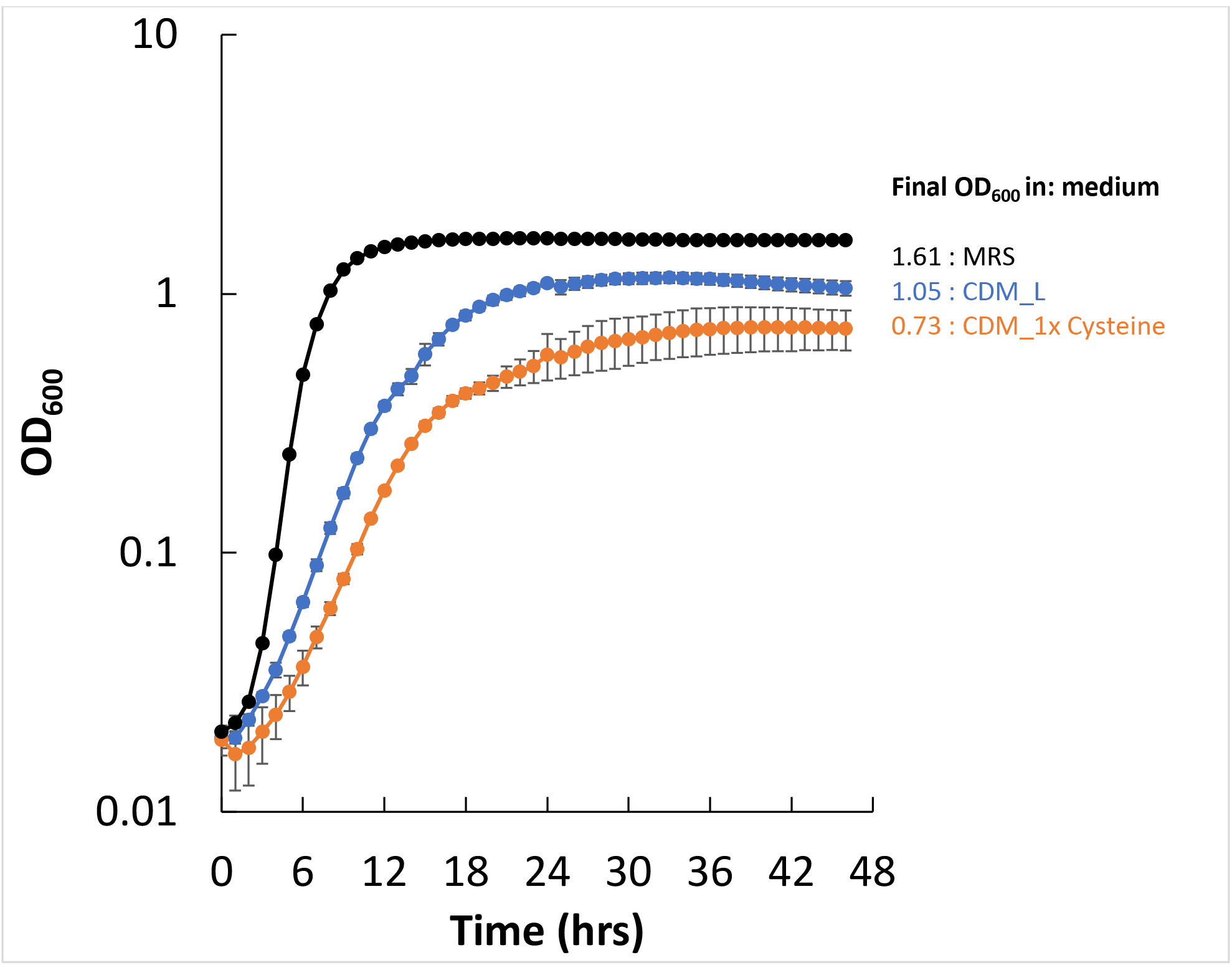
Growth of *Lp. plantarum* NCIMB8826 in MRS, CDM_L, and a CDM without added cysteine. Inoculation at 0.05 from overnight cultures of MRS grown cells washed in PBS. Growth in 96 well plates with 12 technical replicates per growth curve. Note that CDM_L has 10x the cysteine of CDM_1x Cysteine.

**Table S1.** Bacterial strains used in this study.

| Species | Source | Reference |
| --- | --- | --- |
| *Lp. plantarum* WF | Wild *D. melanogaster* isolate | Obadia et al. 2017 |
| *Lp. plantarum* NC8 | Fermented grass | Axelsson et al. 2012 |
| *Lp. plantarum* NCIMB8826 | Human saliva | Hayward & Davis 1956 |
| *Lp. plantarum* LF | Canton-S *D. melanogaster* | Obadia et al. 2017 |
| *Lp. plantarum* endoref | lab *D. melanogaster* | Storelli et al. 2011 |
| *Lp. plantarum* SB001 | lab *D. melanogaster* | Obadia et al. 2017 |
| *Lp. plantarum* SSB | wild *D. melanogaster* | Hardy et al. 2018 |
| *Lp. plantarum* WJL | lab *D. melanogaster* | Ryu et al 2008 |
| *Lp. plantarum* K4 | wheat sourdough starter | Yu et al 2021 |
| *Lp. plantarum* 8.1 | wheat boza | Yu et al 2021 |
| *Lp. plantarum* W1.1 | wheat flour teff injera | Yu et al 2021 |
| *Lp. plantarum* B1.1 | brown flour teff injera | Yu et al 2021 |
| *Lp. plantarum* B1.3 | brown flour teff injera | Yu et al 2021 |
| *Lp. plantarum* T2.5 | fermented tomatoes | Yu et al 2021 |
| *Lp. plantarum* WS1.1 | fermented tomatoes (spoiled) | Yu et al 2021 |
| *Lp. plantarum* 1B1 | cactus fruit (*Opuntia ficus-indicia*) | Tyler et al 2016 |
| *Lp. plantarum* AJ11 | fermented olives | Golomb et al 2013 |
| *Lp. plantarum* BGM55 | fermented olives inoculated with yeast | Golomb et al 2013 |
| *Lp. plantarum* BGM37 | olive fermentation brine | Golomb et al 2013 |
| *Lp. plantarum* BGM40 | fermented olives | Golomb et al 2013 |
| *Lp. plantarum* EL11 | fermented olives | Golomb et al 2013 |
| *Ll. brevis* LF | isolated from lab *D. melanogaster* | Gould et al. 2018 |
| *Ll. brevis* LF2 | isolated from lab *D. melanogaster* | Obadia et al. 2017 |
| *Ll. brevis* SSA | isolated from wild *D. melanogaster* | Hardy et al. 2018 |
| *Ll. brevis* Di | isolated from wild *D. immigrans* | this paper |
| *Acetobacter orientalis* | Canton-S isolate | Gould et al. 2018 |
| *Acetobacter pasteurianus* | Oregon-R isolate | Gould et al. 2018 |
| *Acetobacter tropicalis* | Oregon-R isolate | Gould et al. 2018 |

**Table S2.** Initial growth yield values of strains grown in CDM (Savijoki et al., 2006) with indicated carbon sources. Glu (Glucose), Fru (Fructose), Ac (Acetate), LP (*Lactobacillus plantarum*), (LB) *L. brevis*, (AP) *Acetobacter pasteurianus*, (AT) *A. tropicalis*.

| Final growth yield (OD<sub>595</sub>) |  |  |  |  |  |  |
| --- | --- | --- | --- | --- | --- | --- |
|  | Glu (0.65 %)<br>Fru (0.65 %)<br>Ac (0.1 %) | Glu (0.65 %)<br>Fru (0.65 %) | Glu (1 %)<br>Ac (0.1 %) | Fru (1 %) | Glu (1 %) | Ac (0.1%) |
| LP |  |  | 0.123 |  | 0.056 | 0.001 |
| LB | 0.051 | 0.026 |  |  |  | 0 |
| AP |  |  |  | 0.026 |  | 0.014 |
| AT |  |  | 0.021 |  | 0.044 | 0.021 |

**Table S3.** Variations in CDM and final growth yields of strains. CDM single variations with one component in 10-fold excess. The final growth yield was compared to the final growth yield of the same strains grown in CDM with the final concentration of amino acids, vitamins, nucleotides indicated in Table 1. ODs >0.3 are highlighted. S.E.M. in parentheses.

|  |  | Final growth yield (OD<sub>595</sub>) |  |  |  |
| --- | --- | --- | --- | --- | --- |
|  |  | LP | LB | AP | AT |
| CDM (Table 1) |  | 0.238 (m 0.002) | 0.131 (m 0.001) | 0.103 (m 0.020) | 0.036 (m 0.001) |
| CDM single variations |  |  |  |  |  |
| Ala |  | 0.218 (m 0.009) | 0.138 (m 0.000) | 0.199 (m 0.000) | 0.073 (m0.007) |
| Asn |  | 0.229 (m 0.019) | 0.132 (m 0.005) | 0.136 (m 0.112) | 0.037 (m 0.006) |
| Glu |  | 0.317 (m 0.011) | 0.071 (m 0.006) | 0.030 (m 0.004) | 0.049 (m 0.002) |
| Phe |  | 0.285 (m 0.033) | 0.134 (m 0.002) | 0.062 (m 0.040) | 0.034 (m 0.003) |
| Leu |  | 0.269 (m 0.109) | 0.130 (m 0.002) | 0.025 (m 0.003) | 0.047 (m 0.004) |
| Ile |  | 0.228 (m 0.001) | 0.135 (m 0.011) | 0.064 (m 0.023) | 0.036 (m 0.005) |
| Met |  | 0.231 (m 0.002) | 0.123 (m 0.000) | 0.024 (m 0.041) | 0.040 (m 0.001) |
| Val |  | 0.251 (m 0.044) | 0.138 (m 0.001) | 0.032 (m 0.003) | 0.004 (m 0.001) |
| Trp |  | 0.877 (m 0.535) | 0.127 (m 0.004) | 0.128 (m 0.059) | 0.036 (m 0.003) |
| Gln |  | 0.287 (m 0.007) | 0.076 (m 0.003) | 0.000 (m 0.002) | 0.039 (m 0.000) |
| Arg |  | 0.118 (m 0.030) | 0.124 (m 0.000) | 0.085 (m 0.021) | 0.033 (m 0.002) |
| Gly |  | 0.217 (m 0.005) | 0.133 (m 0.000) | 0.156 (m 0.040) | 0.034 (m 0.000) |
| Lys |  | 0.213 (m 0.000) | 0.141 (m 0.001) | 0.099 (m 0.013) | 0.041 (m 0.000) |
| Pro |  | 0.298 (m 0.016) | 0.155 (m 0.002) | 0.058 (m 0.020) | 0.037 (m 0.000) |
| His |  | 0.268 (m 0.019) | 0.138 (m 0.003) | 0.118 (m 0.083) | 0.040 (m 0.000) |
| Ser |  | 0.296 (m 0.008) | 0.121 (m 0.007) | 0.054 (m 0.002) | 0.065 (m 0.032) |
| Thr |  | 0.257 (m 0.025) | 0.140 (m 0.002) | 0.046 (m 0.011) | 0.032 (m 0.001) |
| Panthothenate |  | 0.246 (m 0.011) | 0.138 (m 0.002) | 0.055 (m 0.027) | 0.027 (m 0.009) |
| Lipoic acid |  | 0.260 (m 0.010) | 0.111 (m 0.004) | 0.032 (m 0.005) | 0.033 (m 0.000) |
| Cyanocobalamin |  | 0.261 (m 0.005) | 0.136 (m 0.000) | 0.088 (m 0.000) | 0.045 (m 0.003) |
| Nicotinic acid |  | 0.218 (m 0.007) | 0.151 (m 0.027) | 0.074 (m 0.002) | 0.039 (m 0.001) |
| para-Aminobenzoic acid |  | 0.217 (m 0.010) | 0.155 (m 0.028) | 0.137 (m 0.065) | 0.038 (m 0.004) |
| Pyridoxine |  | 0.225 (m 0.000) | 0.135 (m 0.001) | 0.067 (m 0.007) | 0.038 (m 0.004) |
| Thiamine |  | 0.231 (m 0.012) | 0.136 (m 0.001) | 0.143 (m 0.051) | 0.037 (m 0.000) |
| Biotin |  | 0.299 (m 0.009) | 0.137 (m 0.005) | 0.075 (m 0.002) | 0.041 (m 0.000) |
| Ascorbate |  | 0.231 (m 0.007) | 0.088 (m 0.052) | 0.316 (m 0.370) | 0.050 (m 0.000) |
| Folate |  | 0.227 (m 0.010) | 0.139 (m 0.001) | 0.071 (m 0.009) | 0.041 (m 0.000) |
| Riboflavin |  | 0.230 (m 0.012) | 0.135 (m 0.000) | 0.187 (m 0.013) | 0.037 (m 0.001) |
| Guanine |  | 0.212 (m 0.004) | 0.136 (m 0.000) | 0.124 (m 0.035) | 0.042 (m 0.005) |
| Uracil |  | 0.267 (m 0.000) | 0.155 (m 0.004) | 0.066 (m 0.019) | 0.043 (m 0.003) |
| Xanthine |  | 0.243 (m 0.007) | 0.138 (m 0.002) | 0.069 (m 0.018) | 0.040 (m 0.003) |
| Adenine |  | 0.252 (m 0.007) | 0.133 (m 0.001) | 0.055 (m 0.033) | 0.044 (m 0.002) |
| PO<sub>4</sub> |  | 0.263 (m 0.008) | 0.208 (m 0.000) | 0.042 (m 0.018) | 0.037 (m 0.002) |
| SO<sub>4</sub> |  | 0.276 (m 0.074) | 0.132 (m 0.000) | 0.120 (m 0.081) | 0.042 (m 0.001) |
| NH<sub>4</sub> |  | 0.200 (m 0.004) | 0.159 (m 0.002) | 0.040 (m 0.017) | 0.036 (m 0.003) |
| Mg |  | 0.205 (m 0.092) | 0.154 (m 0.039) | 0.060 (m 0.003) | 0.039 (m 0.002) |
| Mn |  | 0.290 (m 0.008) | 0.157 (m 0.003) | 0.113 (m 0.029) | 0.057 (m 0.020) |
| Fe |  | 0.210 (m 0.007) | 0.157 (m 0.001) | 0.120 (m 0.029) | 0.042 (m 0.001) |
| Na |  | 0.177 (m 0.032) | 0.139 (m 0.001) | 0.164 (m 0.085) | 0.018 (m 0.000) |
| Bicarbonate, 40 mM |  | 0.347 (m 0.019) | 0.0135 (m 0.001) | 0.022 (m 0.004) | 0.038 (m 0.000) |

**Table S4.**
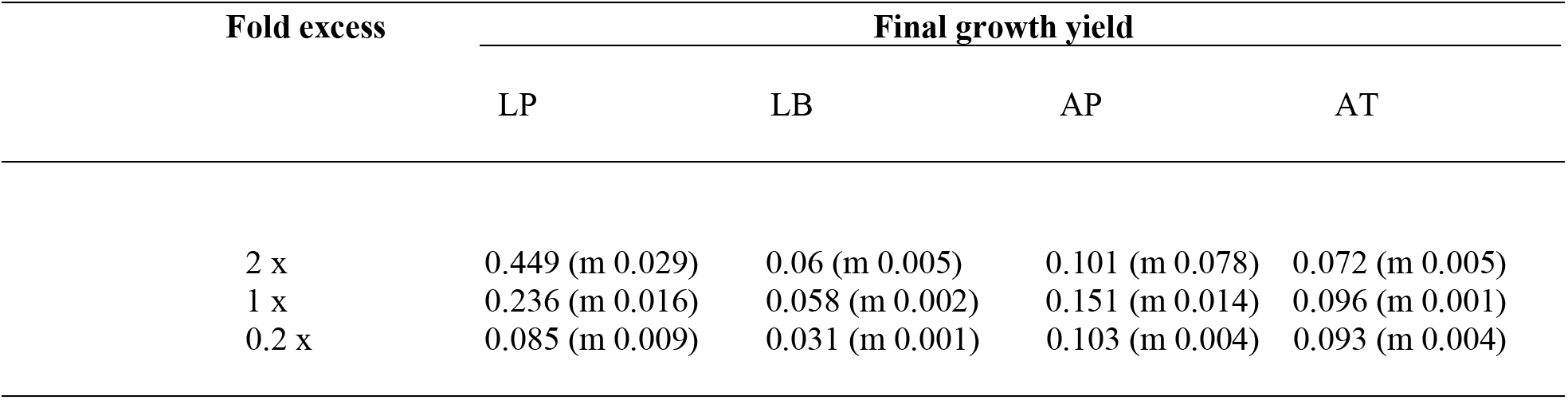
Final growth yield values of strains grown in CDM with indicated fold-difference of the combined amino acids, vitamins, and nucleotides from the concentration of components listed in Table 1, which is set to 1x. LP (*Lactobacillus plantarum*), (LB) *L. brevis*, (AP) *Acetobacter pasteurianus*, (AT) *A. tropicalis*. S.E.M. in parentheses.

**Table S5.** List of chemicals used in this study.

| Chemical | Vendor | Catalog number | Molecular Weight |
| --- | --- | --- | --- |
| MOPS, Free acid, UltraPure | GoldBio | M790 | 209.26 |
| K<sub>2</sub>HPO<sub>4</sub> | Fisher Scientific | P288 | 174.18 |
| NaCl | Fisher Scientific | S271 | 58.44 |
| NH<sub>4</sub>Cl | Fisher Scientific | A649 | 53.49 |
| K<sub>2</sub>SO<sub>4</sub> | Sigma | P9458 | 174.26 |
| MgCl<sub>2</sub>.6H<sub>2</sub>O | Fisher Scientific | BP214 | 203.31 |
| MnCl<sub>2</sub>.4H<sub>2</sub>O | Sigma | M3634 | 197.9 |
| FeSO<sub>4</sub>.7H<sub>2</sub>O | Sigma | F8633 | 278.01 |
| L-Alanine | Sigma | A7627 | 89.09 |
| L-Arginine | Sigma | A5131 | 210.66 |
| Glycine | Sigma | G7126 | 75.07 |
| L-Lysine | Sigma | L5626 | 146.19 |
| L-Proline | Sigma | P0380 | 115.13 |
| L-Histidine | Sigma | H8125 | 155.15 |
| L-Serine | Sigma | S4500 | 105.09 |
| L-Threonine | Sigma | T8625 | 119.12 |
| L-Aspartic acid | Sigma | A9256 | 133.1 |
| L-Asparagine | Sigma | A0884 | 132.12 |
| L-Tyrosine | Sigma | T3754 | 181.19 |
| L-Cysteine-HCl | Sigma | C1276 | 157.62 |
| L-Valine | Sigma | V0500 | 117.15 |
| L-Glutamic acid | Sigma | G1251 | 183.6 |
| L-Tryptophane | Sigma | T0254 | 204.23 |
| L-Phenylalanine | Sigma | P2126 | 165.19 |
| L-Glutamine | Sigma | G3126 | 146.14 |
| L-Leucine | Sigma | L8000 | 131.17 |
| L-Isoleucine | Sigma | I2752 | 131.17 |
| L-Methionine | Sigma | M9625 | 149.21 |
| Ca-(D)-(+)-pantothenate | Sigma | C8731 | 238.27 |
| Lipoic acid | Sigma | T1395 | 206.33 |
| Nicotinic acid | Sigma | 72309 | 123.11 |
| Para-Aminobenzoic acid | Sigma | A9878 | 137.14 |
| Pyridoxine-HCl | Sigma | P9755 | 205.64 |
| Thiamine-HCl | Sigma | T4625 | 337.27 |
| Biotin | Sigma | B4639 | 244.31 |
| Ascorbic acid | Fisher Scientific | AA3623714 | 176.12 |
| Folic acid | Sigma | F7876 | 441.4 |
| Riboflavin | Sigma | R4500 | 376.36 |
| Cyanocobalamin | Sigma | V6629 | 1355.37 |
| Guanine | Sigma | G6779 | 151.13 |
| Uracil | Sigma | U1128 | 112.09 |
| Xanthine | Sigma | X4002 | 152.11 |
| Adenine | Sigma | A2786 | 135.13 |
| Glucose | Sigma | G8270 | 180.16 |
| Fructose | Sigma | F3510 | 180.16 |
| Acetate, sodium salt | Fisher Scientific | S210 | 90.08 |

## Notes

### Competing Interest Statement

The authors have declared no competing interest.

## REFERENCES

Bernhardt C, Zhu X, Schütz D, Fischer M, Bisping B. 2019. Cobalamin is produced by Acetobacter pasteurianus DSM 3509. Appl Microbiol Biotechnol 3875–3885. doi:10.1007/s00253-019-09704-3

Chervaux C, Ehrlich SD, Maguin E. 2000. Physiological Study of Lactobacillus delbrueckii subsp. bulgaricus Strains in a Novel Chemically Defined Medium. Appl Environ Microbiol 66:5306–5311.

Consuegra J, Grenier T, Akherraz H, Rahioui I, Gervais H, da Silva P, Leulier F. 2020. Metabolic cooperation among commensal bacteria supports Drosophila juvenile growth under nutritional stress. ISCIENCE 101232.

da Cruz Pedrozo Miguel MG, Gomes Cardoso P, de Assis Lago L, Freitas Schwan R. 2010. Diversity of bacteria present in milk kefir grains using culture-dependent and culture-independent methods. Food Res Int 43:1523–1528. doi:10.1016/j.foodres.2010.04.031

Degnan PH, Taga ME, Goodman AL. 2014. Vitamin B12 as a Modulator of Gut Microbial Ecology. Cell Metab 20:769–778.

Dysvik A, Leanti La Rosa S, De Rouck G, Rukke E-O, Westereng B, Wicklund T. 2020. Microbial Dynamics in Traditional and Modern Sour Beer. Appl Environ Microbiol 86:1–14.

Elli M, Zink R, Rytz A, Reniero R, Morelli L. 2000. Iron requirement of Lactobacillus spp. in completely chemically defined growth media. J Appl Microbiol 88:695–703.

Grobben GJ, Sikkema J, Smith MR, de Bont JAM. 1995. Production of extracellular polysaccharides by Lactobacillus delbrueckii ssp. bulgaricus NCFB 2772 grown in a chemically defined medium. J Appl Bacteriol 79:103–107. doi:10.1111/j.1365-2672.1995.tb03130.x

Guan H, Yan Y, Li Xiaoling, Li Xiaomei, Shuai Y, Feng G, Ran Q, Cai Y, Li Y, Zhang X. 2018. Microbial communities and natural fermentation of corn silages prepared with farm bunker-silo in Southwest China. Bioresour Technol 265:282–290. doi:10.1016/j.biortech.2018.06.018

Gulitz A, Stadie J, Wenning M, Ehrmann MA, Vogel RF. 2011. The microbial diversity of water kefir. Int J Food Microbiol 151:284–288. doi:10.1016/j.ijfoodmicro.2011.09.016

Hayek SA, Ibrahim SA. 2013. Current limitations and challenges with lactic acid bacteria: a review. Food Nutr Sci 4:73–87.

Henriques SF, Dhakan DB, Serra L, Francisco AP, Carvalho-Santos Z, Baltazar C, Elias AP, Anjos M, Zhang T, Maddocks ODK, Ribeiro C. 2020. Metabolic cross-feeding in imbalanced diets allows gut microbes to improve reproduction and alter host behaviour. Nat Commun 11:4236. doi:10.1038/s41467-020-18049-9

Ho VTT, Fleet GH, Zhao J. 2018. Unravelling the contribution of lactic acid bacteria and acetic acid bacteria to cocoa fermentation using inoculated organisms. Int J Food Microbiol 279:43–56.

Kim G, Huang JH, McMullen JG, Newell PD, Douglas AE. 2018. Physiological responses of insects to microbial fermentation products: Insights from the interactions between Drosophila and acetic acid. J Insect Physiol 106:13–19. doi:10.1016/j.jinsphys.2017.05.005

Li H, Fu J, Hu S, Li Z, Qu J, Wu Z, Chen S. 2021. Comparison of the effects of acetic acid bacteria and lactic acid bacteria on the microbial diversity of and the functional pathways in dough as revealed by high-throughput metagenomics sequencing. Int J Food Microbiol 346:109168. doi:10.1016/j.ijfoodmicro.2021.109168

Pédron T, Mulet C, Dauga C, Frangeul L, Chervaux C, Grompone G, Sansonetti PJ. 2012. A Crypt-Specific Core Microbiota Resides in the Mouse Colon. MBio 3:262–267.

Petry S, Furlan S, Crepeau MJ, Cerning J, Desmazeaud M. 2000. Factors affecting exocellular polysaccharide production by Lactobacillus delbrueckii subsp bulgaricus grown in a chemically defined medium. Appl Environ Microbiol 66:3427–3431.

Ponomarova O, Gabrielli N, Sévin DC, Mülleder M, Zirngibl K, Bulyha K, Andrejev S, Kafkia E, Typas A, Sauer U, Ralser M, Patil KR. 2017. Yeast Creates a Niche for Symbiotic Lactic Acid Bacteria through Nitrogen Overflow. Cell Syst 5:345–357.e6.

Reese V. 1938. Some effects of association and competition on Acetobacter. J Bacteriol 36:357–367.

Ricciardi A, Ianniello RG, Parente E, Zotta T. 2015. Modified chemically defined medium for enhanced respiratory growth of Lactobacillus caseiand Lactobacillus plantarumgroups. J Appl Microbiol 119:776–785.

Rivera-Chávez F, Zhang LF, Faber F, Lopez CA, Byndloss MX, Olsan EE, Xu G, Velazquez EM, Lebrilla CB, Winter SE, Bäumler AJ. 2016. Depletion of Butyrate-Producing Clostridia from the Gut Microbiota Drives an Aerobic Luminal Expansion of Salmonella. CHOM 19:443–454.

Saffarian A, Mulet C, Regnault B, Amiot A, Tran-Van-Nhieu J, Ravel J, Sobhani I, Sansonetti PJ, Pédron T. 2019. Crypt- and Mucosa-Associated Core Microbiotas in Humans and Their Alteration in Colon Cancer Patients. MBio 10:G351–20.

Saguir FM, de Nadra MCM. 2007. Improvement of a Chemically Defined Medium for the Sustained Growth of Lactobacillus plantarum: Nutritional Requirements. Curr Microbiol 54:414–418.

San Roman M, Wagner A. 2018. An enormous potential for niche construction through bacterial cross-feeding in a homogeneous environment. PLoS Comput Biol 14:1–29. doi:10.1371/journal.pcbi.1006340

Sannino DR, Dobson AJ, Edwards K, Angert ER, Buchon N. 2018. The Drosophila melanogaster Gut Microbiota Provisions Thiamine to Its Host. MBio 9.

Savijoki K, Suokko A, Palva A, Varmanen P. 2006. New convenient defined media for [35S]methionine labelling and proteomic analyses of probiotic lactobacilli. Lett Appl Microbiol 42:202–209. doi:10.1111/j.1472-765X.2005.01853.x

Viladomiu M, Hontecillas R, Yuan L, Lu P, Bassaganya-Riera J. 2013. Nutritional protective mechanisms against gut inflammation. J Nutr Biochem.

Wang Z, Shao Y. 2018. Effects of microbial diversity on nitrite concentration in pao cai, a naturally fermented cabbage product from China. Food Microbiol 72:185–192. doi:10.1016/j.fm.2017.12.003

Wang ZM, Lu ZM, Shi JS, Xu ZH. 2016. Exploring flavour-producing core microbiota in multispecies solid-state fermentation of traditional Chinese vinegar. Sci Rep 6:1–10. doi:10.1038/srep26818

Wegkamp A, van Oorschot W, de Vos WM, Smid EJ. 2007. Characterization of the role of para-aminobenzoic acid biosynthesis in folate production by Lactococcus lactis. Appl Environ Microbiol 73:2673–2681.

Wong CNA, Ng P, Douglas AE. 2011. Low-diversity bacterial community in the gut of the fruitfly Drosophila melanogaster. Environ Microbiol 13:1889–1900.

Zheng J, Wittouck S, Salvetti E, Franz CMAP, Harris HMB, Mattarelli P, O’toole PW, Pot B, Vandamme P, Walter J, Watanabe K, Wuyts S, Felis GE, Gänzle MG, Lebeer S. 2020. A taxonomic note on the genus Lactobacillus: Description of 23 novel genera, emended description of the genus Lactobacillus beijerinck 1901, and union of Lactobacillaceae and Leuconostocaceae. Int J Syst Evol Microbiol 70:2782–2858. doi:10.1099/ijsem.0.004107

